# Fungal proliferation before and after the Cretaceous-Paleogene mass extinction in North America

**DOI:** 10.64898/2025.12.11.693767

**Authors:** Rosanna P. Baker, Arturo Casadevall

## Abstract

Palynological evidence of post-catastrophe fungal proliferation after global calamities has been found for the Permian-Triassic and Cretaceous-Paleogene (K/Pg) extinction events. However, unlike the globally documented post-Permian fungal bloom, evidence of a post-Cretaceous event has previously been limited to a single site in New Zealand. Our analysis of a K/Pg boundary section from the Denver Basin in Colorado revealed a fungal proliferative spike occurring immediately after the Chicxulub impact. The discovery of a post-impact fungal bloom in North America corroborates the New Zealand finding and supports the interpretation that this was a global phenomenon. We also identified a prolonged interval of elevated fungal abundance in the Late Cretaceous, dating to approximately 30,000-10,000 years before the impact, temporally correlated to a period of climatic cooling at the site and intriguingly coincident with the Poladpur phase of the Deccan Traps. Taken together with reports of fungal expansion following prior global calamities, these findings indicate that fungi can often flourish in the aftermath of ecosystem-level collapse. Given the capacity of fungi to cause disease in both plants and animals, the occurrence of fungal proliferative events has major potential implications for the recovery of surviving species after global cataclysms.

**Significance Statement:** Fungal proliferation evidenced by spikes in fungal abundance in geologic samples can signify major ecosystem disruptions. Such spikes are well documented globally for the Permian-Triassic extinction but for the Cretaceous-Paleogene extinction have been reported previously only in New Zealand. Here, we describe a North American fungal bloom that occurred immediately after the Chicxulub impact, as well as an earlier fungal spike in the Late Cretaceous that coincided with a cooling event hypothesized to have been driven by Deccan volcanism. Identification of this pre-impact fungal proliferative episode provides additional support for the uranium-lead-based model of Deccan eruptive rates, which places the high-volume Poladpur pulse tens of thousands of years before the bolide impact.

## Introduction

The concept of disaster microbiology posits that major disturbances to macroscopic ecology by calamities are paralleled by disturbances in environmental microbiology (1, 2). This phenomenon is prominent in the fossil record where large spikes in fungal abundance have been documented following the mass extinction events at the ends of the Permian (3) and Cretaceous periods (4). Fungal proliferation after the Permian extinction is documented for diverse geographic sites in the supercontinent of the Pangea (3) but for the Cretaceous-Paleogene (K/Pg) boundary, has only been reported from a single site in New Zealand (4). Although direct microfossil evidence of post-K/Pg fungal proliferation has not been reported in North America, indirect support was provided by the detection of the amino acid α-aminoisobutyric acid, a fungal product, in the boundary strata of the Raton and Power River basins (5).

Fungi are the great degraders of organic matter in the biosphere. Hence, calamities that result in mass death events provide food for fungal growth thereby providing a mechanistic explanation for the proliferation of fungi in the post-cataclysm environment. However, mass mortality may not be required for fungal proliferation since ecological upheavals can also weaken resistance of extant species to fungal diseases. In this regard, surviving species may have increased susceptibility to fungal diseases because of weakened immunity caused by environmental stress, nutritional deficits, and habitat degradation, which combined with larger inocula from proliferating fungi, could make them susceptible to mycoses. Notably, fungi typically prefer cooler temperatures and acidic environments (6–8), and geologic calamities that result in planetary cooling, reduced sunlight, and acid rain could promote the fungal growth.

Understanding the microbiological disturbances that accompany past mass-extinction events is important because that knowledge can provide additional insight into the scope of the disaster and its effects on surviving species. Fungi are major pathogens of both plants and animals and any fungal proliferation following a cataclysm has the potential to profoundly affect the health of surviving biota, which may be more vulnerable to mycotic diseases from the environmental disruption. In this regard, it was noted that fungal microfossils at the end of the Permian resemble current plant pathogenic fungi raising the possibility that surviving life would also have to deal with a fungal threat (3). Similarly, fungal proliferation at the end of the Cretaceous has been proposed to have contributed to the rise of mammals, since these are disproportionately resistant to mycotic diseases relative to ectothermic vertebrates such as reptiles (9).

In this study we have revisited the question of whether fungi proliferated on a global scale following the K/Pg mass extinction event using samples from North America. Our results confirm a fungal spike at the K/Pg boundary which supports the hypothesis that this mass extinction, like the one marking the end of the Permian, was followed by a worldwide interval of increased fungal activity. In addition, we report a prolonged fungal bloom that occurred tens of thousands of years before the bolide collision at a time when Deccan volcanism was producing a cooler, more acidic climate that would be expected to favor fungal growth.

## Results

### Study areas

Sediment samples were collected from three well-documented and stratigraphically constrained K/Pg boundaries, each with various primary markers (e.g., boundary clay, shocked minerals, iridium anomaly) of the Chicxulub impact: the Bowring Pit from the West Bijou study area (D1 Sequence) in the Denver Basin, Colorado (10–13) and the Mud Buttes (14) and John’s Nose sections (15, 16), both from the Williston Basin, North Dakota (Fig. 1). The samples analyzed span the latest Cretaceous, K/Pg boundary clay, and earliest Paleocene strata.

**Fig. 1.**
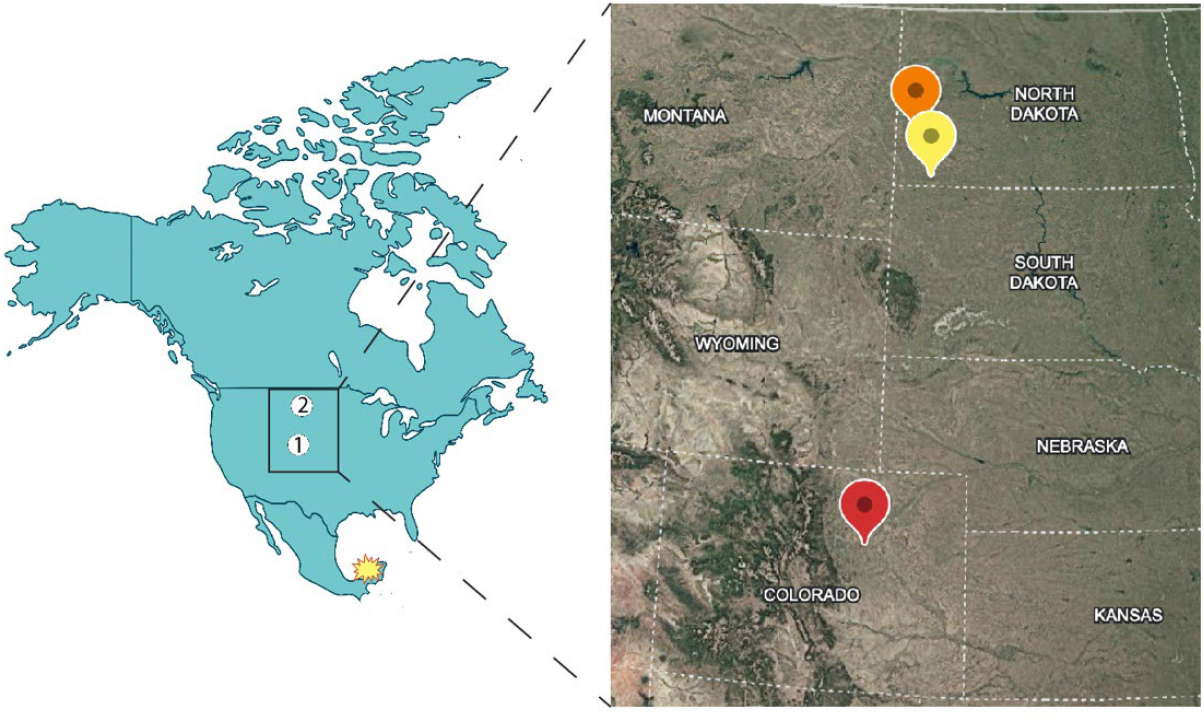
Map of North America (left panel) showing the study areas in Colorado (A) and North Dakota (2) in relation to the site of the Chicxulub impact (yellow) and magnification of the study areas (right panel): Bowring Pit from the West Bijou study area (D1 Sequence) in the Denver Basin, Colorado (red), Mud Buttes (yellow) and John’s Nose (orange) from the Williston Basin, North Dakota. The distances of the Colorado and North Dakota locations from the bolide impact site in the Yucatan are approximately 2500 and 5500 km, respectively. Prepared using BioRender and Google Earth.

### Evidence for periods of fungal proliferation in the Bowring Pit

The palynology of the Denver Basin, and in particular, the Bowring Pit at West Bijou has been extensively documented (11, 17, 18). These studies detailed the genera and relative abundances of plant species before and after the boundary in North American K/Pg formations, established palynomorph biozones, and documented the abrupt extinction of numerous palynomorphs but did not identify the K/Pg fungal bloom that was reported more than two decades ago in New Zealand (4). We focused our analysis to determine the proportion of microfossils that were fungal in origin versus palynomorphs and tailored our methods to maximize the retention and detection of fungal forms. Most of the samples analyzed in the high-resolution Bowring Pit section were proportionally higher in palynomorphs, except for two distinct layers where fungal forms constituted 50% or more of the total assemblage (Fig 2A). The fungal proliferation we identified at the K/Pg boundary established that a mass extinction-associated fungal bloom occurred not only in New Zealand, but also in North America. Unexpectedly, we also observed an extensive period of fungal predominance in Late Cretaceous strata that predate the boundary by tens of thousands of years (Fig 2A). Spike samples, ranging from ∼50-60% fungal, were found in lignite strata but most other lignite samples in the section were less than 30% fungal (Fig 2A), which demonstrated the discriminatory power of our method. The two periods of fungal proliferation in the Bowring Pit section were distinct in the assemblage of fungal morphotypes visualized. The most abundant morphotypes in the K/Pg fungal spike were fungal hyphae and large, dicellate ellipsoidal spores (Fig 2B) but several other forms were observed, ranging from small globose and ovoid spores to large unicellular and multicellular forms. The Late Cretaceous fungal bloom was comparable in diversity to the K/Pg spike but predominantly consisted of small, dark brown globose and ovoid spores (Fig 2C).

**Fig. 2.**
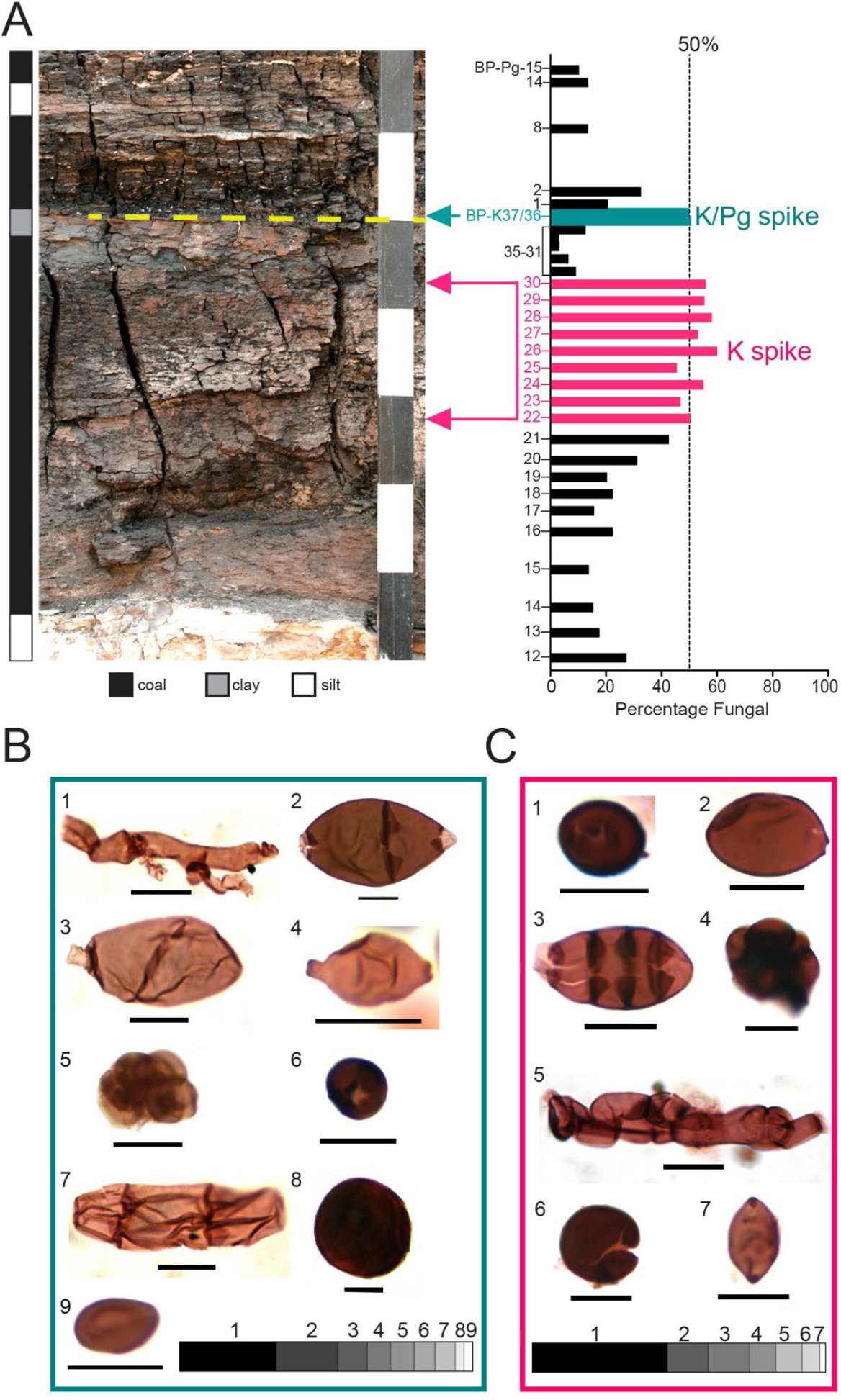
Fungal spikes in the Bowring Pit section. (A) Photograph on left showing lithostratigraphy with K/Pg boundary indicated by a dashed yellow line (scale = 10 cm) aligned with bar graph (right panel) of the percentage fungal forms among the total microfossil count in each sample. Fungal percentages were approximately 50% in a spike at the boundary (K/Pg spike, turquoise) and for an extended period below the K/Pg boundary (K spike, pink). (B) The K/Pg spike consisted of one-third hyphae (1) and two-thirds spores that were 21% 25-44 µm dicellate ellipsoidal (2), 10% 11-22 µm unicellate diporate (3), 8% 21-35 µm unicellate ellipsoidal (4), 8% 24-70 µm multicellate globose (5), 7% 5-11 µm unicellate globose (6), 7% 28-40 µm multicellate elongated (7), 3% 24-70 µm unicellate globose (8), and 3% 9-14 µm unicellate ovoid (9). (C) Unicellate globose spores 4-12 µm in diameter (1) were the most abundant morphotype in the K-spike, followed by 14% 9-18 µm unicellate ovoid (2), 14% 16-60 µm multicellate ellipsoidal (3), 9% 11-56 µm multicellate globose (4), 9% hyphal fragments (5), 6% 13-25 µm unicellate globose (6), and 2% 9-18 µm unicellate diporate (7). The slide designation and England Finder coordinates for each image in B and C are reported in Table 1 and scale bars = 10 µm. Photograph in panel A was provided by Tyler Lyson and is credited to Rick Wicker, both at the Denver Museum of Nature & Science, (Denver, CO.)

### Cretaceous and Paleogene fungal spikes in North Dakota

Our analysis of strata from the Mud Buttes and John’s Nose sections in North Dakota each revealed two spikes in fungal proliferation that occurred before and after the K/Pg boundary (Fig. 3). The post K/Pg spike at Mud Buttes occurs in one sample collected at the top of a lignite at the base of the Fort Union Formation, 4 cm above the base of the K/Pg boundary clay, and above the fern spike reported from this section which occurs in the base of the lignite (14). The underlying boundary clay and overlying fine-grained mudstone unit lack fungal spikes. The post K/Pg spike at Mud Buttes, like the K/Pg spike in the Bowring Pit, was approximately one-third fungal hyphae but the remaining two-thirds differed in morphotype composition and included multicellate and unicellate globose spores as well as dicellate and multicellate ellipsoidal spores (Fig. 3B, upper panel). The Cretaceous spike occurs in two samples, ∼4 cm below the boundary clay in a fine-grained mudstone paleosol. Samples above and below are from the same facies unit and lack a fungal proliferation. This fungal spike was the most uniform of the spikes analyzed in this study and consisted almost entirely of dark brown globose spores that ranged in size from 13-22 µm in diameter and occasionally occurred in 2-4 cell clusters (Fig. 3B, lower panel).

**Table 1.** Locations of fossils in representative images from Figures 2 and 3.

| **Fungal spike** | **Morphotype description** | **Slide name** | **EF location** |
| --- | --- | --- | --- |
| Bowring Pit K/Pg | (1) hypha | 070825_BP-K-36_1A | L382 |
|  | (2) dicellate ellipsoidal | 070825_BP-K-37_1A | P400 |
|  | (3) unicellate diporate | 070825_BP-K-36_2A | L403 |
|  | (4) unicellate ellipsoidal | 070825_BP-K-36_1A | O472 |
|  | (5) multicellate globose | 070825_BP-K-37_1A | L420 |
|  | (6) unicellate globose | 070825_BP-K-37_1A | L352 |
|  | (7) multicellate elongated | 070825_BP-K-36_1A | J412 |
|  | (8) unicellate globose | 070825_BP-K-37_2B | N81 |
|  | (9) unicellate ovoid | 070825_BP-K-36_1A | M453 |
| Bowring Pit K | (1) unicellate globose | 081225_BP-K-27_1A | M500 |
|  | (2) unicellate ovoid | 081225_BP-K-30_1A | G363 |
|  | (3) multicellate ellipsoidal | 081925_BP-K-22_1A | X440 |
|  | (4) multicellate globose | 081925_BP-K-24_1A | P410 |
|  | (5) hypha | 081225_BP-K-28_1A | N363 |
|  | (6) unicellate globose | 081925_BP-K-24_1A | L460 |
|  | (7) unicellate diporate | 081225_BP-K-28_1A | O440 |
| Mud Buttes Pg | (1) hyphal | 022725_FU4_2A | O414 |
|  | (2) multicellate globose | 022725_FU4_2A | U404 |
|  | (3) unicellate globose | 022725_FU4_2B | O100 |
|  | (4) dicellate ellipsoidal | 022725_FU4_2B | N133 |
|  | (5) multicellate ellipsoidal | 022725_FU4_2A | U464 |
| Mud Buttes K | (1) unicellate globose | 060425_HC12_3A | T433 |
|  | (2) 2-cell cluster | 032125_HC13_1B | G153 |
|  | (2) 3-cell cluster | 032125_HC13_1A | K414 |
|  | (2) 4-cell cluster | 060425_HC12_2A | P373 |
| John's Nose Pg | (1) unicellate globose | 072225_TRL48_1A | H373 |
|  | (2) hypha | 072225_TRL48_1A | N390 |
|  | (3) unicellate ovoid | 072225_TRL48_1A | N430 |
|  | (4) dicellate ellipsoidal | 072225_TRL48_1B | F94 |
|  | (5) multicellate globose | 072225_TRL48_1A | M374 |
|  | (6) multicellate elongated | 072225_TRL48_1B | P111 |
| John's Nose K | (1) hypha | 090925_TRL34_1A | L421 |
|  | (2) multicellate globose | 090925_TRL34_1A | Q440 |
|  | (3) multicellate ellipsoidal | 090925_TRL30_1A | L334 |
|  | (4) unicellate ovoid | 090925_TRL30_1A | J370 |
|  | (5) multicellate elongated | 090925_TRL30_1A | N450 |
|  | (6) unicellate globose | 090925_TRL34_1A | N334 |
|  | (7) unicellate globose | 090925_TRL30_1A | K371 |

**Fig. 3.**
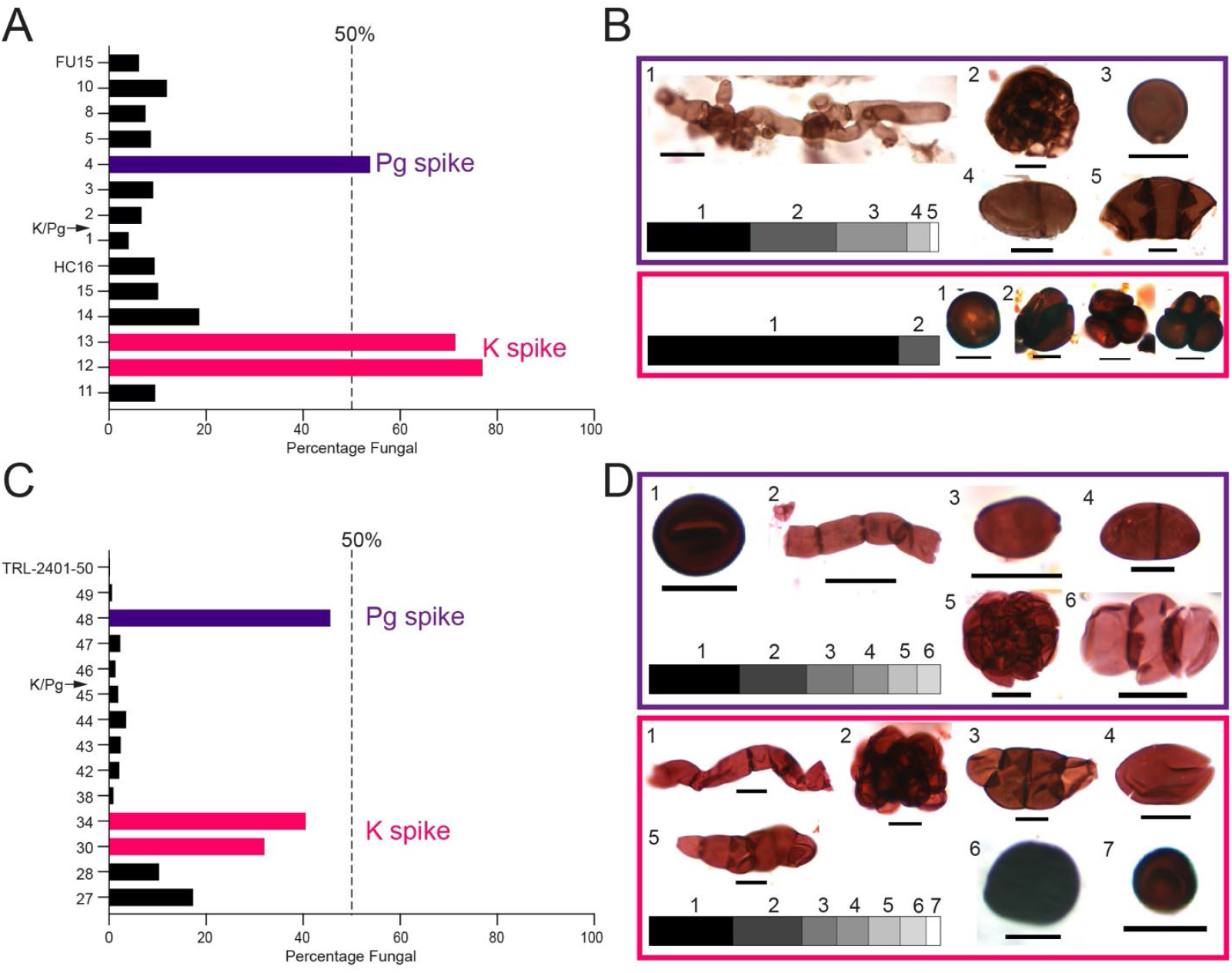
Fungal spikes in the Mud Buttes and John’s Nose sections (Cedar Creek Anticline, Williston Basin, North Dakota). (A) Bar graph of the percentage of fungal microfossils in each Mud Buttes sample showing fungal proliferative spikes above (Pg spike, purple) and below (K spike, pink) the K/Pg boundary. (B) Representative images of fungal morphotypes in the Mud Buttes Pg spike (upper panel, purple) that were 35% hyphal (1), 30% 20-34 µm multicellate globose (2), 24% 9-14 µm unicellate globose (3), 8% 20-23 µm dicellate ellipsoidal (4), and 3% 29-53 µm multicellate ellipsoidal (5) and K spike (lower panel, pink) that consisted of 86% 13-22 µm unicellate globose spores (1) and 14% 2-4 cell clusters ranging in size from 21-24 µm (2). (C) Bar graph of the percentage of fungal microfossils in each John’s Nose sample showing spikes in fungal proportions above (Pg spike, purple) and below (K spike, pink) the K/Pg boundary. (D) The John’s Nose Pg spike (upper panel, purple) was composed of 28% 7-12 µm unicellate globose spores (1), 23% hyphae (2), 16% 8-18 µm unicellate ovoid (3), 12% 18-23 µm dicellate ellipsoidal (4), 10% 13-25 µm multicellate globose (5), 8% 15-39 µm multicellate elongated (6), and 3% 13-14 µm unicellate globose (7) and the K spike (lower panel, pink) consisted of 29% hyphae (1), and several spore types including 24% 14-44 µm multicellate globose (2), 12% 33-47 µm multicellate ellipsoidal (3), 11% 12-20 µm unicellate ovoid, 11% 23-83 µm multicellate elongated (5), 9% 14-45 µm unicellate globose (6) and 4% 7-8 µm unicellate globose (7). The slide designation and England Finder coordinates for each image are reported in Table 1. Scale bar = 10 µm.

The fungal spikes observed in the John’s Nose section were smaller in proportion than those in the other two sites, reaching approximately 40% of the fossilized assemblage. The Late Cretaceous spike occurs in a blocky carbonaceous unit 22-29 cm below the boundary clay. The early Paleocene spike occurs in a lignite, 4 cm above the boundary clay, and above the reported fern spike from this section. Both the Late Cretaceous and early Paleocene spikes were rich in fungal hyphae but most of the fungal spores were larger in the former compared to the latter (Fig. 3D). The early Paleocene spikes in the two North Dakota sections occur after the fern spike, whereas the fungal blooms in the Bowring Pit and New Zealand section preceded the proliferation of ferns.

### Fine sieving depletes samples of fungal microfossils

We extended our qualitative analysis of fungal morphotypes by conducting a quantitative assessment of the fungal spore sizes in each of the six proliferative spikes (Fig 4A). The mean spore size in most of the fungal spikes ranged from approximately 12-17 µm except for the Bowring Pit K/Pg and John’s Nose K spikes that had average spore sizes of 29 µm and 26 µm, respectively (Fig 4A). Whereas most spores in these two fungal spikes were larger than 20 µm, most spores in the other four spikes were smaller than 20 µm.

**Fig. 4.**
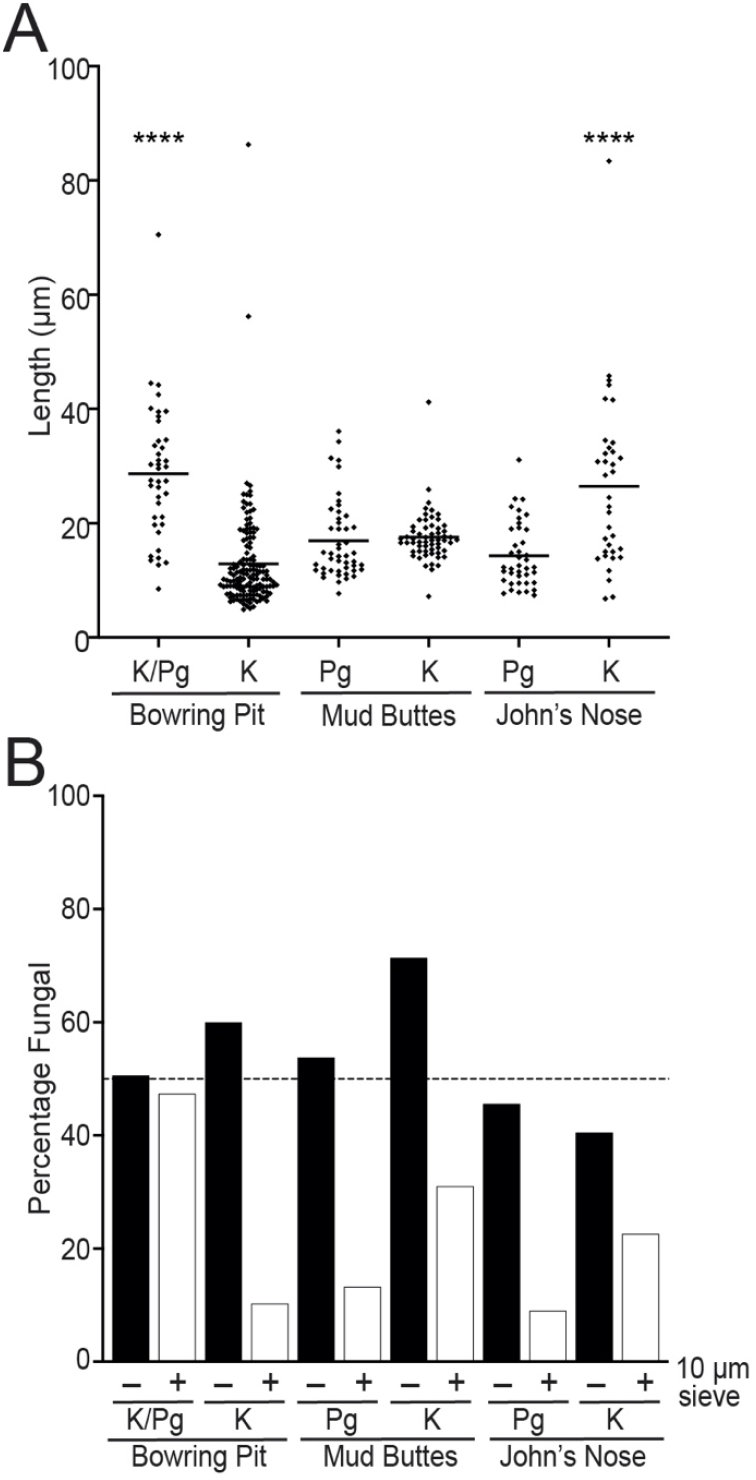
Effect of fine sieving on fungal fossil retention. (A) Scatter plot comparing lengths fungal spores shows significantly larger average spore sizes for Bowring Pit K/Pg and John’s Nose K spikes when analyzed for statistical significance by one-way analysis of variance (ANOVA) with Tukey’s multiple comparisons test, ****p < 0.0001. Bars denote mean length. (B) Comparison of the percentage of fungal spores retained in samples without (–) and with (+) 10 µm sieving. Fungal percentages decreased dramatically for most samples after fine sieving with the smallest differences observed for the Bowring Pit K/Pg and John’s Nose K spikes that had the largest average spore sizes.

As our method omitted the final 10 µm sieving step that is routinely used in palynological preparations, we wondered how this fine sieving step would impact the retention of fungal spores in our samples. We compared percentages of fungal spores for samples prepared without and with 10 µm sieving and noted that the four spike samples with small average spore sizes showed a marked depletion of fungal spores after fine sieving (Fig 3B) that resulted in a shift in fungal percentages from above to below 50%. The John’s Nose K spike, which had larger average spore sizes, decreased from 40% to 23% fungal (Fig 4B). Only the Bowring Pit K/Pg spike that had an average spore size of 29 µm was unaffected, remaining approximately 50% fungal after fine sieving (Fig 4B). This analysis implies that five of the six fungal spikes reported here would not have been identified in these samples if they were prepared according to traditional methods.

### The Bowring Pit Cretaceous fungal bloom coincided with the Deccan Poladpur phase

In addition to the expected post-impact spike, we observed a prolonged interval of elevated fungal abundance in the Late Cretaceous strata of the Bowring Pit section. Time calibration using dated ash layers (13) allowed us to estimate the timing of this Cretaceous spike to approximately 22,000-10,000 years before the Chicxulub impact (Fig 5). Notably, this coincides with a transient ∼5°C decrease in mean annual air temperature recorded at the site (19) (Fig 5). We also identified Late Cretaceous fungal spikes in the two North Dakota sections, although the lack of precise time-calibration data precludes assignment of definitive ages to those events.

**Fig. 5.**
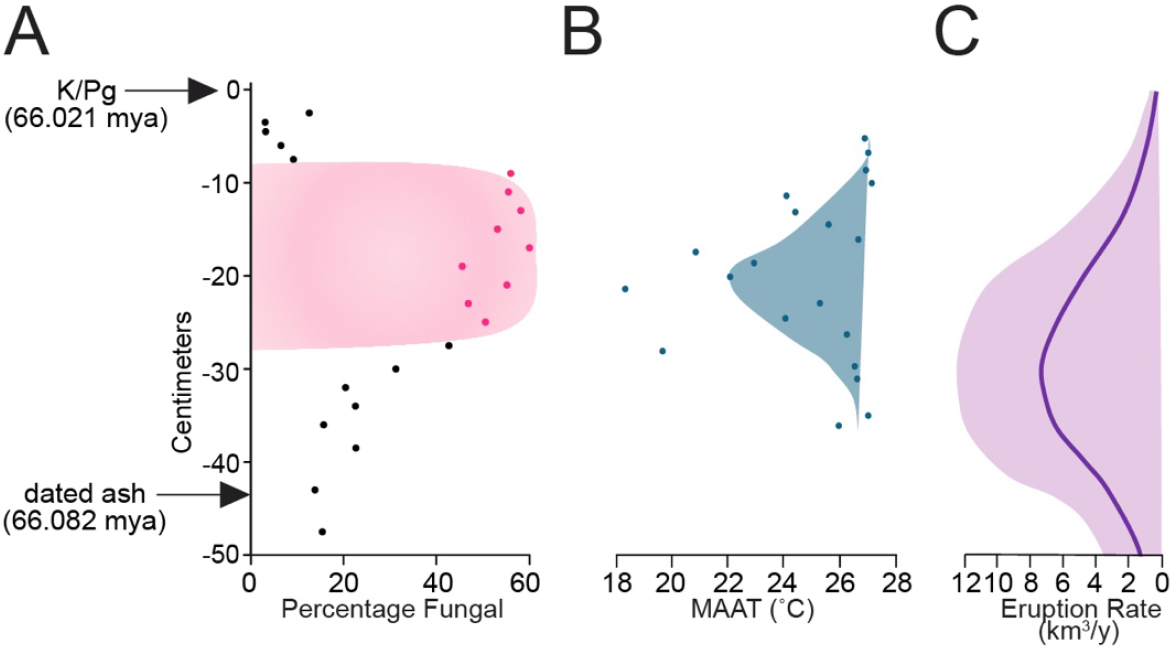
Pre-boundary fungal proliferation coincides with Deccan-driven cooling. (A) Time-calibration data from the Bowring Pit section (13) places the Cretaceous fungal spike at approximately 10-30 kyr before the K/Pg extinction event. (B) A transient ∼5°C decrease in mean annual air temperatures (MAAT) was measured at the Bowring Pit section in the same time interval (19). (C) The high-volume Deccan Poladpur pulse is estimated to have occurred 50-10 ky before the K/Pg boundary based on the uranium-lead eruption rate model (37). B and C are redrawn representations of published data (19, 37).

## Discussion

Evidence for fungal proliferation following the K/Pg mass extinction was previously documented in New Zealand (4) but has not been corroborated in other geographic regions in the intervening two decades, although detection of a fungus-produced amino acid in K/Pg sediments provided indirect evidence for fungal proliferation in the northern interior of the United States (5). Our identification of a prominent fungal spike at the K/Pg boundary in the high-resolution Bowring Pit section of the Denver Basin in Colorado suggests that a global fungal bloom occurred in the immediate aftermath of the Chicxulub impact.

Standard palynological procedures vary among studies but typically involve demineralization with hydrochloric and hydrofluoric acids, oxidation, alkali treatment, and acetolysis (20). Whereas pollen grains readily withstand these processes, recovery of non-pollen palynomorphs, including fungal spores, can be improved by the use of gentler methods (21, 22). To maximize the preservation of fungal microfossils, we employed treatment with sodium hexametaphosphate, a non-acid deflocculant that is particularly effective for clay-rich sediments because its charged phosphate ions adsorb onto clay particles and disaggregate them (23). We also omitted the common 10 µm sieving step, which removes amorphous organic material, but also risks the loss of small fungal spores. In addition to the Bowring Pit K/Pg fungal spike, we detected blooms of fungal proliferation in the Late Cretaceous in this section and both above and below the boundary at the two sites in North Dakota. Although the fungal spikes in early Paleocene strata in the two North Dakota sites do not correlate temporally with known global catastrophes, we note that all three sites showed concordance in each having early Paleogene and Late Cretaceous fungal spikes. Importantly, the fungal morphotype assemblages of the Bowring Pit K/Pg spike, which contains nearly three-quarters large spores and hyphal fragments, differs markedly from the other fungal spikes identified in this study that are dominated by small globose to ovoid spores. Assigning the Bowring Pit K/Pg morphotypes to specific extant fungal genera is beyond the scope of this study, but it is likely that the abundant decaying biomass in the wake of mass extinction favored the proliferation of saprophytic forms. We were unable to match the fungal morphotypes identified at the three sites with extant species, which together with the fact that fungal forms were restricted to discrete layers, argues against contamination. A taxonomic analysis of the Cretaceous and Paleogene morphotypes will be the subject of future studies.

An increased prevalence of fungi in post-disaster ecosystems could be expected to produce both beneficial and detrimental effects. Fungi may have offered the benefit of an alternative source of nutrition in landscapes where plant life had been severely diminished. Notably, fungal agriculture in ants has been dated to approximately 66 million years ago, roughly coincident with the fungal blooms described here, when the high prevalence of mycotic biomass could have selected for an adaptation to cope with widespread food scarcity (24). Conversely, pathogenic fungi could have imposed additional stress on organisms already struggling to survive. Although the fossilized forms identified here cannot be confidently assigned to extant animal pathogenic lineages based on morphology, the possibility of fungal pathogenicity in the K/Pg fungal bloom warrants further investigation. In this regard, large inocula could compensate for low intrinsic pathogenicity of a given species and still cause disease (25). Thus, fungal proliferation before and after the K/Pg extinction, together with its associated pathogenic potential, would lend support to the fungal infection mammalian selection hypothesis that posits a selective advantage for mammals over reptiles owing to thermal restriction and advanced immunity of mammalian hosts (9).

The Late Cretaceous fungal bloom observed in the Bowring Pit aligns well with the Poladpur phase of Deccan volcanism which was associated with global cooling (19, 26). Deccan-driven climate cooling, coupled with increased soil acidity resulting from chemical fallout, would be expected to favor fungal proliferation, as most genera grow optimally below 30°C and within a pH range of 5-6 (6–8). Consistent with this expectation, an abundance of mycorrhizal fungi accompanied by a marked decrease in pollen spores was documented in late Maastrichian strata from the Chhindwara region and Nand-Dongargaon basin of the Deccan volcanic province (27). Identifying a North American fungal bloom concurrent with Deccan trap activity suggests that a major global ecosystem disruption preceded the Chicxulub impact and lends further support to the hypothesis that Deccan volcanism contributed to the K/Pg mass extinction event (28).

Our findings, together with the earlier report of a fungal bloom at the K/Pg boundary in New Zealand (4) argue that the K/Pg mass extinction event, like the end-Permian, was followed by a period of global fungal proliferation and implicate a broader capacity for fungal spore abundance in palynological records to serve as an indicator of catastrophic environmental disruptions. In addition to the K/Pg bloom, we identified an earlier fungal spike predating the Chicxulub impact that may reflect ecological stress associated with Deccan volcanism. Other than the primarily impact-driven K/Pg extinction, the other four Phanerozoic mass extinctions have been linked to massive volcanic eruptions as evidenced by the temporal link between large igneous provinces and extinctions at the end-Ordovician (29), late Devonian (30), end-Permian (31), and end-Triassic (32). A recent study demonstrating a strong correlation between bulk eruption rate and extinction magnitude argues that the Deccan Traps emissions were large enough in magnitude to have played a significant role in the K/Pg extinction (33). Indeed, high-latitude palynological records indicate a gradual phase of decline preceding the K/Pg boundary, consistent with prolonged environmental stresses induced by Deccan activity (34–36). Our identification of a pre-boundary fungal spike supports a “two-hit’ scenario of K/Pg extinction, in which Deccan volcanism initiated an early decline that rendered many taxa more vulnerable to extinction after the impact. Finally, we note that examination of sedimentary deposits for evidence of fungal proliferation may provide a valuable tool for identifying additional periods of ecological stress throughout geologic history, including events that were regional rather than global in extent, with the caveat that recovery of fungal microfossils is highly influenced by sample preparation and filtration parameters.

## Materials and Methods

### Non-acid extraction protocol

Palynomorphs were processed according to a modified method for non-acid extraction described previously (23). Sediment samples were crushed into grain size using a mortar and pestle, heated at 60°C for 1 h in 3% Alcojet, then incubated overnight at room temperature. Sodium hexametaphosphate was added at 1:2 (w/w) and samples were heated at 60°C, with stirring, for 40 min. Following coarse sieving, the > 500 µm fraction was centrifuged in a saturated solution of sodium metatungstate at 2,500 *g* for 5 min. The concentrated organic fraction was washed with distilled water and stained for 5 min with Gram’s safranin solution or an additional fine-sieving step with a 10 µm filter was included prior to staining. Strew mounts were prepared by embedding in 4% polyvinyl alcohol and mounting on slides using Eukitt Quick-hardening mounting medium.

### Microscopic analysis

Slides were examined using an Olympus AX70 microscope under bright field illumination with 40X or oil-immersion 100X objectives and QCapture-Pro 6.0 software was used to capture images with a Retiga 1300 digital charge-coupled device camera. Microfossil positions on slides were recorded using an England Finder (Table 1). Fungal forms were identified based on morphology, resemblance to other fossilized fungal spores and extant fungal species, and melanization of cell walls, features that distinguish them from pollen palynomorphs. Fungal spore length measurements in pixels were made from 40X images using ImageJ software and measurements were converted to micrometers using the conversion factor of 0.0167µm/pixel.

## Acknowledgments

We are grateful to Drs. Mary Berbee and Antoine Bercovici for insightful discussions. We are indebted to Dr. Tyler Lyson for providing the soil samples, for guiding us through the stratigraphy, and for making many helpful suggestions in the preparation of this manuscript. A.C. was supported in part by National Institutes of Health R01 grants HL059842, AI152078, and AI052733.

